# Developing A Programmable, Self-Assembling *Squash Leaf Curl China Virus* (SLCCNV) Capsid Proteins Into “Nano-Cargo”-Like Architecture: A Next-Generation “Nanotool” For Biomedical Applications

**DOI:** 10.1101/338269

**Authors:** Raja Muthuramalingam Thangavelu, Deepan Sundarajan, S.U Mohammed Riyaz, Michael Immanuel Jesse Denison, Dharanivasan Gunasekaran, Rajendran Ganapathi, Nallusamy Duraisamy, Kathiravan Krishnan

## Abstract

A new era has begun in which pathogens have become useful scaffolds for nanotechnology applications. In this research/study, an attempt has been made to generate an empty cargo-like architecture from a high-profile plant pathogen of *Squash leaf curl China virus* (SLCCNV). In this approach, SLCCNV coat protein monomers are obtained efficiently by using a yeast *Pichia pastoris* expression system. Further, dialysis of purified SLCCNV-CP monomers against various pH strengthenened (5–10) disassembly and assembly buffers produced a self-assembled “Nanocargo”-like architecture, which also exhibited an ability to encapsulate the magnetic nanoparticles at *in vitro*. Bioinformatics tools were also utilized to predict the possible self-assembly kinetics and bioconjugation sites as well. The biocompatibility of “SLCNNV-CP-Nanocargo” particles was also evaluated by *in vitro* cancer cells, which eventually proved the particles to be versatile material for the next generation “nanotool” capable of housing various therapeutic or imaging agents.

## Introduction

There has always been a high demand for biological entities which have the ability to act as a multifaceted scaffold or template. Biological entities having intrinsic self-assembling properties are recognized as an excellent aspect in nanotechnology applications [1-4]. The coat protein of viruses are the kind of self-assembling peptides which form a fine architecture at a nanoscale precision is called capsid. Recently viruses are being studied for their ability to self-assemble into the nanoscale particles of discrete size and definite geometry [5-7]. These self-assembling features of viruses have been used as nanoscaffolds for a wide range of innovative applications in various fields such as sensing, imaging, drug delivery, biocatalysis and bioelectronics [8]. Viral nanotechnology is a newly emerging multi-disciplinary area, has begun to manipulate hundreds of both mammalian and plant pathogenic viruses towards the next-generation smart nanodevices [9,10]. The reason of sudden emergence of virus nanotechnology is the search for the solutions to several fundamental challenges in nanoparticles fabrication, that is, controlling the traits of synthetic nanoparticles such as monodispersity, assembly, stability, size and morphology [11]. Viruses derive portions of their structures from peptide-based complexes called ‘capsids’. These complexes form nanoscale structures that are often self-assembling, monodispersing, symmetric and stable under general physiological conditions [12, 13]. These tiny structures, also described as viral nano-particles (VNPs) by materials science researchers, possess numerous traits that make them outstanding candidates for nanoresearch. Another superior choice for biomedical researchers is virus-like particles (VLPs), a non-infectious counterpart from plant viruses. These are particularly advantageous because they are less likely to be pathogenic in humans and therefore less likely to induce undesirable side effects [8]. They can be easily produced in sufficient quantities with recombinant technologies. In addition, their properties are easily programmable through changes at the genome level to produce novel functionalities[14].The VLPs that are currently being developed for biomedical applications share the common property of being self-assembling – they form a closed capsid-like structure with an altered symmetryfrom a limited number of protein subunits, which are also referred to by various terms with the prefix of “nano”-cage, -carrier, -cargo, and -container[8]. This states that an interior environment is capable of housing, therapeutic or imaging agents and the exterior environment is capable of multivalent presenting for cell targeting [4,15,16]. There are many studies that have reviewed the self-assembling properties of various kinds of animal and plant virus particles under various *in vitro* conditions such as temperature, pH, and ionic strength [5, 17-22]. In particular, three plant viruses have been studied more than hundred times by various researchers across the country specifically for nanotechnology applications: *Cowpea mosaic virus, Cowpea chlorotic mottle virus* and *Tobacco mosaic virus* [8, 23]. Though a few species of viruses have been studied extensively for nanotechnology purposes, there is a need for a new kind of virus that is superior to existing ones. Indeed, exploring novel VNPs or VLPs and their self-assembling properties will lead to novel ideas and concepts for fabrication for human welfare. With this aim, we chose a well-known plant pathogenic virus of the *Begomovirus* isolates of *Squash leaf curl China virus* (SLCCNV) for this study, whose molecular pathogenicity we had already established in our laboratory studies. The true reason for our choice of this virus is no study yet to done to reveal the structural and assembly properties of *Begomoviruses*, which are also crucial for coat-host protein interaction mediated pathogenicity studies.

The *Squash leaf curl China virus* (SLCCNV), a plant pathogenic virus, is one of the 200 species in the *Begomoviruses* genus and belongs to the *Geminiviridae* family [24]*. Begomoviruses* are responsible for the huge loss of many economically important crops such as tomatoes, chilly, cassava, squash and cotton [24, 25].The genome of *Squash leaf curl China virus* (SLCCNV) and other members of this genus generally consists of a single-stranded (SS) DNA molecule (5422 bp in size). It encodes eight open reading frames, where the AV1 gene (771bp) of the coat protein (CP), represents a twinned icosahedral capsid morphology which encapsulates the whole nucleic acid genome [26]. Interestingly, all members of the *Geminiviridae* have the unique capsid morphology of geminate particles (non-enveloped), consisting of two incomplete T=1 icosahedral joined together to produce a structure with 22 pentameric capsomers, made of 110 identical CP subunits, each consisting of approximately 30 kDa (256 AA) with an isoelectric point (pI) of 10 [27, 28]. The size of the virus particle is ∼38 nm in length and 22 nm in diameter.

This research work aimed to unveil the self-assembling properties of the native SLCCNV particles as well as *Pichia* expressed SLCCNV coat protein subunits towards. We chose the assembly buffer medium as a reflection of the biological conditions in the intracellular environment [29,30]. To support this experimental study, a detailed *in silico* analysis was also performed to predict the SLCCNV coat protein structure, assembly and its conjugation sites. [31-33]. In vitro A549 cell line studies also demonstrated biocompatible properties of SLCCNV coat protein. A successful *in vitro* self-assembly is valid only if it can be exploited to selectively entrap materials within the mechanisms of the SLCCNV-CP-assembly. By encapsulation with magnetic nanoparticles (MNPs) it was also confirmed [34]. MNPs were used in this study because of their wide range of application ranging from medical diagnosis to treatment [35;36].

## Materials and methods

### Materials

All the chemicals for growth media and supplements were purchased from Himedia Laboratories, India. Dialysis membrane (15kDa cut-off) also purchased from Himedia Laboratories, India. All the glasswares (Borosil, Scott Duran) and chemicals for buffers and other components for reagents including uranyl acetate and sodium phosphotungstic acid were obtained from Sisco Research Laboratories, India. *E. coli* strains of DH5 alpha, GS115 *Pichia pastoris* strain and pPICZαA plasmid (Invitrogen, USA) for cloning were kind gifts from Prof. S. Meenakshi Sundaram, Centre for Biotechnology, Anna University, Chennai, India. The antibiotic Zeocin was purchased from Invitrogen, USA. For western blotting, the primary polyclonal antibody of DSMZ *African cassava mosaic virus* (ACMV) was used, which was a kind gift from Dr. V. G. Malathi, Distinguished Professor and Virologist, Tamil Nadu Agriculture University, Coimbatore, India. Goat anti-rabbit secondary antibody conjugated HRP enzyme and coat protein-specific forward and reverse primers were obtained from Genei Technologies, Bengaluru. Polyvinylidenedifluoride (PVDF), cloning enzymes and plasmid isolation kits were purchased from Thermo Fischer and Fermentas, Hyderabad. The TAS-ELISA kit for ACMV was purchased from DSMZ, Germany. For chromatography, pre-packed HiTrap ion exchange columns were purchased from GE Healthcare, India. For cell line studies, A549 lung cancer lines were obtained from the NCCS cell repository, Pune. Acetaminophen, potassium bromide and MTT [3-(4,5-dimethylthiazol-2,5-diphenyltetrazolium bromide] were obtained from Sigma-Aldrich, India. Ham’s F12-K medium and TPVG were purchased from Himedia, India. Foetal bovine serum (FBS) and Antibiotic-Antimycotic was from Gibco Life Technologies, USA. Ferrous chloride hydrated extra pure (code no. 03846) and ferric chloride anhydrous 98% extra pure (code no. 03817) were obtained from LobaChemie Pvt. Ltd, India.

### Isolation and purification of virus particles

To investigate the assembling and disassembling processes, native *Squash leaf curl China virus* (SLCCNV) was isolated from the natural host *Benincasa hispida* (winter melon), which was collected from a field at Perambalur in the southern part of the state of Tamil Nadu in India. Further, screenings and confirmations were made by molecular studies including PCR amplification with coat protein-specific primers, which was already reported in our previous work [37]. The virus particles or so-called viral nanoparticles (VNPs) were purified by cesiumsulfate (Cs_2_SO_4_) density gradient ultracentrifugation (CP100WX, HITACHI, JAPAN) using fixed angle rotor P100AT2 (803,000xg) and swing bucket rotor P55ST2 (366,000 x g) with necessary modifications [38]. Purified fractions of virus proteins were collected and dialyzed against 0.1M phosphate buffer (pH 7.2) to remove the remnants of Cs_2_SO_4_. The purity of the native virus particles was also confirmed through SDS-PAGE followed by a western blot analysis [39,40]. For experimental control, the purified native virus particles were used all along with SLCCNV-CP subunits to be expressed in yeast *Pichia pastoris*.

### Heterologous expression of SLCCNV-CP in yeast *Pichia pastoris*

The coat protein expression comprises three principal steps: (a) insertion of the gene into an expression vector; (b) transformation of the expression vector into the *P. pastoris*host; (c) examination of potential strains for the expression of the SLCCNV-CP gene. The methylotrophic yeast *Pichia pastoris* GS115 strain was applied as a host cell to express SLCCNV coat protein [41]. The plasmid pPICZαA containing methanol-inducible AOX1 promoter was used to construct the full length SLCCNV coat protein gene [42]. The major advantage of using pPICZαA (3.6 kb) vector for expression is the presence of a Zeocin-resistant gene for selection, which has an alpha-factor secretion signal for directing the secreted expression of the recombinant protein into the growth medium [43].

### Construction of expression vector

To design an insert for expression, a complete coat protein gene (771bp) from the AV1 region of DNA-A of SLCCNV was taken from our own nucleotide deposit in the NCBI repository (GenBank accession no.**KF188433.1**) [37]. The sense and anti-sense primers KKCPF-5’-CCGGAATTCATGGCGAAGCGACCACCACCAGATA-3’ and KKCPR-5’-CCGGGTACCAT TTGTTACCGAATCCATAAAA-3’ capable of generating a 771-bp coat protein gene fragment (256 amino acids) containing double restriction enzyme sites (*Eco*RI and *Kpn*I) were designed using the BioEdit 7.1 software [44]. To express the pure form of SLCCNV coat protein without the C-terminal peptide, stop codon (5’ cap, *poly*-A tail) had been included in the anti-sense primer. By PCR amplification, the double-stranded oligonucleotide fragments corresponding to a CP protein of SLCCNV, with *Eco*RI and *Kpn*I cohesive ends, were obtained by annealing (63˚C-65˚C) the sense and anti-sense primers using the whole SLCCNV genome as a template. The PCR-obtained CP gene fragments of SLCCNV were inserted into multiple cloning sites (MCS) of pPICZαA vector in the C-terminal in-frame fusion. In brief, the inserts of the CP gene and pPICZαA vectors were appropriately digested with restriction enzymes *Eco*RI and *Kpn*I. An enzyme-excised fragment was inserted between the *Eco*RI and *Kpn*I sites of the pPICZαA vector, yielding clone plasmids of pPICZαA-SLCCNV-CP. The final construct was confirmed by restriction mapping with the same digestive enzymes. Following this, a propagation of vector was performed by the transformation of those clones into *E. coli* DH5-α-competent cells derived from calcium chloride, which was screened and then cultured in a low-salt LB medium containing the Zeocin antibiotic (25μg/ml). Furthermore, all expression steps were performed according to the manual given by Easyselect™ *Pichia* expression kit (Invitrogen) with necessary modifications.

### Transformation, screening and confirmation of Mut^+^ phenotype

For *P. pastoris* transformation, the host cells were washed in two simultaneous steps with Milli-Q water and 1M sorbitol, based on manufacturer guidelines. The 100μl of the fresh, competent GS115 *P. pastoris* cells plus 10μg of linearized (*Sac*I enzyme digested) plasmid DNA was pulsed in 0.1 cm electroporation cuvettes (Model.No.620, BTX, Harvard apparatus, Holliston, USA) at 1.5 kV, 50 F, 250 Ω and 10–12 millisecond (Bio-Rad gene-pulser).The transformed cells were selected on YPD (Yeast extract peptone dextrose) Zeocin plates and then screened for the insert by PCR on isolated yeast genomic DNA using 5’ and 3’AOX1 (5’-GACTGGT TCCAATTGA CAAGC-3’and 5’-GTCCCTATTTCAATCAATTGAA-3’). Screening of recombinants was confirm with SLCCNV coat protein, specific primers at 771bp. Subsequently, positive clones were used for the analysis of the methanol-utilizing phenotype. The Mut+ phenotype for the transformed GS115 cells was determined by growing clones on a minimal methanol medium with histidine (MMH) and a minimal dextrose medium with histidine (MDH) plates. A single transformed colony of GS115 was grown in 200ml of BMGY (Buffered Glycerol-Complex Medium) at 28–30˚C and at the agitation rate of 250 RPM. The cells (OD600 = 2-6) were harvested and resuspended in 500 ml of BMMY (Buffered Methanol-Complex medium) in a 1L triple-side baffled flask (OD600 = 1.0) to induce expression at 28–30˚C. Absolute methanol was added to a final concentration of 0.5% every 24 hours to maintain induction for up to four days. Each day, 1ml of the supernatant was collected to optimize the protein expression profile of transforming Mut^+^ phenotype cells. The collected supernatant was subjected to electrophoretic separation on a 12% polyacrylamide gel (SDS-PAGE) and stained by silver nitrate according to the standard procedure [39]. Following electrophoresis, proteins were electrotransferred onto a PVDF membrane and immunoblotted with an ACMV primary antibody specific to the *Begomovirus* coat protein. The secondary HRP enzyme conjugated antibody used to detect the protein of interest at the end [40]. The concentrated supernatant of GS115 transformed with an empty vector (pPICZαA) was used as the negative control in both tests. The percentage of *Pichia*-expressed SLCCNV-CP was then determined by ELISA with the known concentration of the ultracentrifuge-purified native virus protein as a positive control [40].

### Purification and characterization of expressed SLCCNV-CP

At the end of the fourth day of culture in BMMY, the supernatant was harvested by high-speed cooling centrifugation (10000 RPM), after it was concentrated (10-to 100-fold) by the usage of 75% ammonium sulphate precipitation [46]. The precipitant was further dialyzed against a 0.1M phosphate buffer (pH 7.2) to eliminate the ammonium salts. Then the concentration of protein was determined by absorbance measurement at 280nm (1cm path length) with an extinction coefficient of 3.4. The ion exchange chromatography (IEC) purification was performed based on the chromato-focussing method. This method was chosen to separate the SLCCNV-CP according to the pI (isoelectric point) and its theoretically determined pKa values [17]. To purify the coat protein from the active ammonium sulfate fraction, one millilitre of the fraction was loaded into the ACTA purifier (GE Healthcare) in 5ml of HiTrap ion exchange column. The column was pre-equilibrated with buffer A and buffer B. The unbound proteins were washed with the same buffer and the bound proteins were eluted by a linear gradient of salt (1M NaCl) with a flow rate of 1 ml/min. The eluted fractions containing the SLCCNV-CP were pooled and stored at −80˚C. The obtained fractions were analyzed at 12% SDS-PAGE. Following electrophoresis, SDS-PAGE bands containing the protein of interest were excised from stained gels and subjected to the in-gel trypsin digestion procedure as described elsewhere [47, 48]. Spectral measurements were taken using an Ultraflex III MALDI-TOF/TOF instrument (Bruker Daltonics, Germany)in the positive ion reflector mode for acquiring the peptide mass fingerprint (PMF) at the International Center for Genetic Engineering and Biotechnology (ICGEB), New Delhi, India. Tandem mass spectrometry-based fragmentation was employed to identify the protein from the observed peptide precursor ions. The instrument parameters for PMF (peptide mass fingerprinting) and MS/MS analysis were set as described elsewhere [48]. The fragmented peptides were analyzed using flexAnalysis software (version 3.0) and sent to database search using BioTools software (version 3.2). The database search parameters were set as described [48] and fragment masses with MS tolerance of up to +/-100 ppm and MS/MS tolerance of up to +/-0.75 Da were searched in the NCBInr database. The following criteria were used for identification: (i) significance threshold was set to achieve p<0.05, (ii) the expectancy cut-off was set to 0.05, and (iii) individual ion score ≥ 45 was only considered for identification.

### Investigation of self-assembly of SLCCNV (native and *Pichia*-expressed)

*In vitro* assembly were performed by the conventional dialysis method (15kDa cut-off membrane) against various pH assembly buffers. The assembly buffers with different pH (5 – 10) were prepared by the addition of 1mM CaCl_2_ [29,30,49, 50]. In the first study, small volume of native SLCCNV particles presence in the buffer of pH 7.2 (0.1M phosphate buffer) were dialysis against the buffer of pH 5.0 (0.1M sodium acetate) at the cold temperature for 8 hours. After that, the existing buffers in dialysis tank were exchanged to the other assembly pH buffers of pH of 6.0 - 8.0 (0.1M sodium phosphate), pH 9.0 (0.1M sodium borate) and pH 10.0 (0.1M sodium glycine) at the stipulated time period of 8 hours in cold conditions. Eventually, the same dialysis procedure were carried out for *Pichia*-expressed SLCCNV-CP subunits. Meanwhile small aliquots of the dialyzed samples from the different assembly pH buffers were taken to the study by High-resolution transmission electron microscopy (HRTEM-FEI, TechnaiG2, 30S-TWIN D905, USA) and dynamic light scattering (DLS, Malvern Instruments Ltd., Malvern, UK). For the HRTEM analysis, the specimens were processed with a 2% negative stain of sodium phosphotungstate and uranyl acetate prepared at neutral pH. From the HRTEM micrograph, it was observed that the sodium phosphotungstate-stained samples were clearer than the uranyl acetate-stained samples [51].

### *In silico* analysis of SLCCNV coat protein

Initially, a converted FASTA format of a coat protein-related amino acid sequence (accession no. **KF188433.1**)[37] was submitted to the I-TASSER (Iterative Threading Assembly Refinement) server [31,52]. Kind of similar protein structure templates were threaded through the PDB library by LOMETS, a locally installed meta-threading approach. The excised PDB templates were reassembled into full-length models using Monte Carlo simulations, with the threading unaligned regions built by *ab initio* modelling. When no appropriate templates were identified by LOMETS, the whole protein structures were built by *ab initio* modelling. SPICKER identified low free energy state through clustering of the simulation decoys. Finally, TMalign was used to align the LOMETS templates and the PDB structures. The final full atomic models were obtained from the I-TASSER decoys by REMO [45]. Further, it was validated through UCLA-DOE (http://nihserver.mbi.ucla.edu/Verify-3D/) structure evaluation server, which gives a visual analysis of the quality of a putative crystal structure for proteins. The structural models were validated using PROCHECK [45]. The query-modelled protein structure was submitted as PDB files on the SAVES server (http://nihserver.mbi.ucla.edu/SAVES/) [53,54]. The quality of the protein structure was validated from the Ramachandran plot through the Procheck server [55].

### Prediction of assembly of SLCCNV-CP monomers

The PatchDock server is a simple and intuitive web interface, available at http://bioinfo3d.cs.tau.ac.il/PatchDock [56]. The I-TASSER-derived monomeric structure of SLCCNV coat protein was uploaded to the server and a docking request was submitted. Once the prediction process was completed, the results were received by email containing the web link for the predicted page. Modelling was performed to assemble a pentameric coat protein structure and the it was repeated at three times. At the end of the process, pentameric structure of the SLCCNV capsid protein was received. In addition, we tried to predict the solvent accessible surface areas (SASA) by Accelrys Discovery Studio 2.5, because it was believed that one of the key factors involved in the coat protein subunit assembly. Therefore, each modelled protein structure (monomer to pentameric) was uploaded in Accelrys Discovery Studio 2.5 [57]. The Connelly-type solvent accessibility helped in the identification of buried and exposed residues in the solvent system. The solvent-exposed and buried residues were displayed as red and blue colour, respectively.

### Analysis of binding sites and surface functional groups in the SLCCNV-CP

Molsoft ICMPro is a very useful bioinformatics tool in the analysis of protein structure. The tools included in the ICM Pro are calculating RMSD, identifying closed cavities, calculating contact and surface area, measuring angles, distances and generating Ramachandran plots [58]. The pentameric protein structure was loaded into the Molsoft ICM-Pro molecular software for the analysis of binding sites and surface groups. The protein structure was loaded into the software window and converted into an ICM object for further process. The ICM-converted protein structure was used for the prediction of binding sites and the residues on the surface of the protein. With the same bioinformatics tool, the possible bioconjugation sites were also identified in the ICM object. The predicted bioconjugation sites were indicated as molecular surfaces in the 3D ribbon diagram of SLCCNV coat protein.

### Biocompatibility test

General biocompatibility or toxicity tests, aimed mainly at detection of the biological activity of test substances, can be carried out in many cell types (e.g. fibroblasts, HeLa, hepatoma cells) [59]. In this research/study, the biocompatibility feature of the expressed SLCCNV-CP alone was determined in a dose-dependent manner by the MTT assay with A549 lung cancer cell line. [60]. The A549 cells (1×104) were seeded onto 96 well flat-bottom culture plates and incubated for 24 hours at 37˚C with 5% of CO_2_ to reach the confluence in 70%. Over thirty various concentrations of SLCCNV-CP (0.54 to18 μg/ml) were tested against the cell culture seeded in the 96 well plate. The treated cells were kept incubated for 24 hours, after which 10µL of MTT (5mg/ml in PBS) added to each well and the plate was incubated for another four hours at 37˚C. The resulted formazan crystal was dissolved in 100µL of DMSO (Dimethyl sulfoxide) with a gentle shake and the absorbance was measured at 595nm using an ELISA reader (BioTek power wave XS, USA). All these steps were replicated at three times independently and the average has been shown as a cell viability percentage in comparison with the control experiments, where the SLCCNV-CP untreated controls were considered as 100% viable.

Cytotoxicity%=[1-(OD _treated_/OD _control_)]*100

where OD_treated_ and OD_control_ corresponded to the optical densities of treated and control cells. The percentage of cytotoxicity (%) plotted as a function of concentration, fitted to a sigmoidal curve and the median inhibitory dose (IC_50_) value was determined on the basis of this curve. The IC_50_ represents the concentration of *Pichia*-expressed SLCCNV-CP required to achieve 50% inhibition of cell growth at *in vitro*.

### Encapsulation efficiency of SLCCNV-CP-Nanocargo on magnetic nanoparticles

To evaluate the encapsulation efficiency of SLCCNV-CP-Nanocargo, negatively charged magnetic nanoparticles (MNPs) with the mean size of ∼25nm were used. The magnetic nanoparticles (MNPs) were prepared using the co-precipitation method followed by a slight modification of protocol, which has been reported by us earlier [61]. To assemble the SLCCNV-CP over MNP core, one equivalent of MNPs (about 5.2μg/ml) was mixed with 100 equivalents of SLCCNV-CP monomers and first dialyzed against a pH 6 disassembly buffer at 4˚C for overnight (∼12 hours). Subsequently, the samples were collected and freshly dialyzed against an assembly buffer of pH 7 at 4˚C in order to re-assemble the SLCCNV-Nanocargo with MNPs. Finally, the dialyzed samples were pooled and centrifuge at 50,000g for 1hour at 4˚C and then the pellet containing the samples was resuspended in 100μl of a 0.1M phosphate buffer (pH 7.2). The samples were stored at −20˚C until use. The encapsulation efficiency of SLCCNV-CP was determined by UV-Vis spectroscopy, HRTEM coupled with energy dispersive X-ray spectroscopy (HRTEM-FEI, TechnaiG2, 30S-TWIN D905, USA) and 8% native polyacrylamide gel electrophoresis.

## Results

### SLCCNV-CP expressed by yeast *Pichia pastoris*

The expression of the entire *Begomovirus* unmodified coat protein subunits was first achieved in the eukaryotic yeast *Pichia pastoris* system. The PCR-amplified product corresponding to the full-length coat protein gene of 771bp is shown in the agarose gel image (Figure 1A). The double-enzyme (*Kpn*I & *Eco*RI) digested restriction map of the expression cassette of pPICZαA-SLCCNV-CP was reviewed which provides the product length of 3.6 kb for the pPICZαA vector and 771bp for the CP gene, whereas undigested gives 4.4 kb (Figure 1B). The cloned expression vectors were successfully propagated into recA1-and endA1-deficient *E. coli* DH5-α strains with good yield. The DNA sequencing results of AOX1 5’ and 3’ primer-amplified PCR products determined that our recombinant construct had an insert in the correct orientation (sequencing data not included). An enzyme (*Sac*I)-linearized vector of pPICZαA-SLCCNV-CP (4.4kb) was successfully electrotransferred to GS115 cells, which yielded Zeocin-resistant *Pichia* recombinants that were screened and confirmed by culturing them with Zeocin antibiotics (Figure 1C&D). In figure 1E,the gel image shows PCR-amplified products at 771bp and 500bp with CP gene-specific primers confirming the presence of the CP gene in the genome of *Pichia* recombinant colonies. The recombinants cultured in BMMY growth medium yielded the SLCCNV-CP protein by secretion, which was collected by centrifugation and confirmed in the protein denaturing gel. In figure 1F, silver stained gel image of SDS-PAGE, lane 1 shown the multiple protein bands for the supernatant collected from the transformed colonies and the lane 2 dedicated to the non-transformed colonies shown only few bands. An another SDS-PAGE performed with both purified native SLCCNV particles (Figure 1G lane1) and *Pichia* expressed CP monomers (Figure 1G lane 2) has shown some equivalency at the high molecular weight bands. The purified expressed SLCCNV-CP monomers in the gel, shown multiple bands of different molecular weight and one at 30kDa corresponding to the MW of SLCCNV-CP monomer (Figure 1G lane 2). In the denaturing gel with native virus particles (Figure 1G lane1), presence of high MW bands might represent the stable capsomeric architecture (Virus/110monomers/30kDa=3,300kDa). We surprised of the stability of the native virus particles against common protein denaturing conditions. Even at a twofold increased concentration of reducing agents (β-mercaptoethanol and Dithiothreitol) and SDS, the native virus particles remains intact and not yield any monomeric protein bands in the gel. Finally, the intact native virus particles denatured at 10 min incubation in high temperature (85°–90˚C). It has given two denatured bands with a high molecular weight close to one another. Meanwhile, to confirm the native virus stability in the temperature, a Differential Scanning Calorimetry (DSC) was performed [62]. The DSC result indicated that the whole virus particles were intact and stable up to 75˚C. It was only started to denature at the temperature above 80˚C (Figure S1, supplementary information), thus added one more evidence of the stability of virus particles.

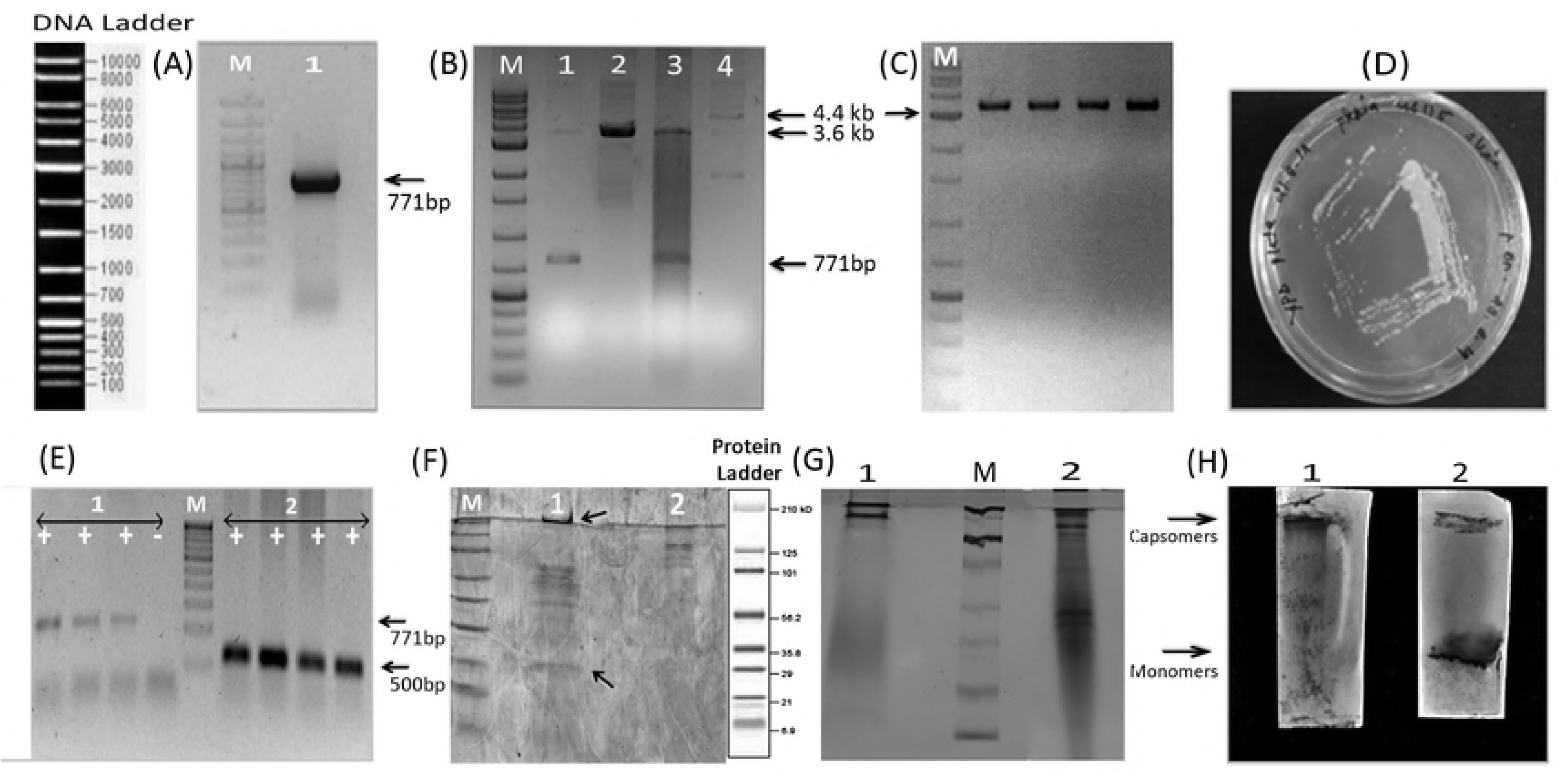

This phenomenon was also confirmed after performing the confirmative assay of western blot using an ACMV polyclonal antibody and an HRP-conjugated secondary antibody with DAB (3, 3’-diaminobenzidine) substrate. In the Figure 1H, the blotting membrane showed the bands corresponding to the protein samples similar in the previous SDS-PAGE gel picture (Figure 1G). A single intensified band that is exactly the same size observed in the SDS-PAGE for native virus particles observed in the Figure 1H Panel 1. Another western blot (Figure 1H Panel 2) with *Pichia*-expressed SLCCNV-CP showed two high-intensity bands roughly equivalent to coat protein monomers in 30kDa, and multimers of capsomers which were similarly observed in SDS-PAGE. The high yield of SLCCNV coat protein monomers was achieved by inducing the culture of *Pichia* recombinants with 0.5% methanol at different time points (4 days), which yielded a threefold increase in the overall concentration of protein on the last day (compared to the first day), as quantified by ELISA (Fig.S2., supplementary data). The untransformed cultures of *Pichia* cells (control) in the BMMY medium decreased in growth (OD_600_ at 3 to 1) in the presence of methanol from the second day itself. It shows that untransformed colonies are much more susceptible to methanol due to the absence of a methanol-utilizing gene/vector.

### Ion exchange chromatography and MALDI-TOF

The active fraction from the ammonium sulphate fractionation was loaded into a weak cationic ion exchange chromatography [CM-Sepharose Fast Flow 5ml column]. Initially, the column was pre-equilibrated with buffer A (20 mM Tris, pH 6.0, containing 150mM NaCl). The protein was unabsorbed by the column [Fig.S3.,Supplementary information(inset CM-ion exchange profile image)]. After this, the unbound protein from the CM -Sepharose was adjusted to pH 7.8 and loaded onto a strong anionic Qff 5ml column. Unbound proteins were collected as flow-through with Buffer A, [20mM pH 7.8 containing 150mM NaCl about 6 CV (column volume)] and the bound proteins were eluted at about 12 CV by Buffer B (20mM Tris pH 7.8 +1M NaCl) using a linear gradient method. Figure S3., (Qff-ion exchange profile image) shows the elution profile of the active fraction. It is clear that flow-through proteins do not contain the protein whereas the elution fractions show the purified protein. Homogeneity and purity of the elution fraction was determined through 12% SDS-PAGE (fig.S3. insetSDS-PAGE image). In SDS-PAGE, a similar banding pattern of multiple protein bands was observed at a different molecular weight that represents the monomeric (30kDa) and multimeric forms of coat protein. These multimers can either be dimer, trimer, tetramer, pentamer or hexamer. In addition, high MW protein bands were excised and utilized for the MALDI analysis. The ELISA quantified the final concentration of the purified *Pichia*-expressed SLCCNV coat protein monomers was 0.57mg/ml which also twofold higher than the ultracentrifuge purified native virus protein (0.241mg/ml) (Figure S2). The protein sequence information obtained from MALDI-TOF/TOF of a single excised coat protein spot was submitted to the MASCOT search engine (Matrix Science Ltd.). The MASCOT results showing the list of all entries from the NCBInr for known protein sequences matched the experimental peptide sequence data (Figure 2). It revealed that PMF (protein mass fingerprint) obtained from MALDI-TOF/TOF analysis from the 1D gel was identical to the *Squash leaf curl China virus* and the protein coverage was achieved at 39%. Twelve peptides were matched with the *Begomovirus* coat protein sequence profile through an analysis using the NCBI database. The protein scores achieved greater than 74. The predicted molecular mass (30kDa) and pI (10) value of the expressed SLCCNV-CP were found to be identical with the *Begomovirus* isolates and particularly the *Squash leaf curl China virus* of the Thailand strain.

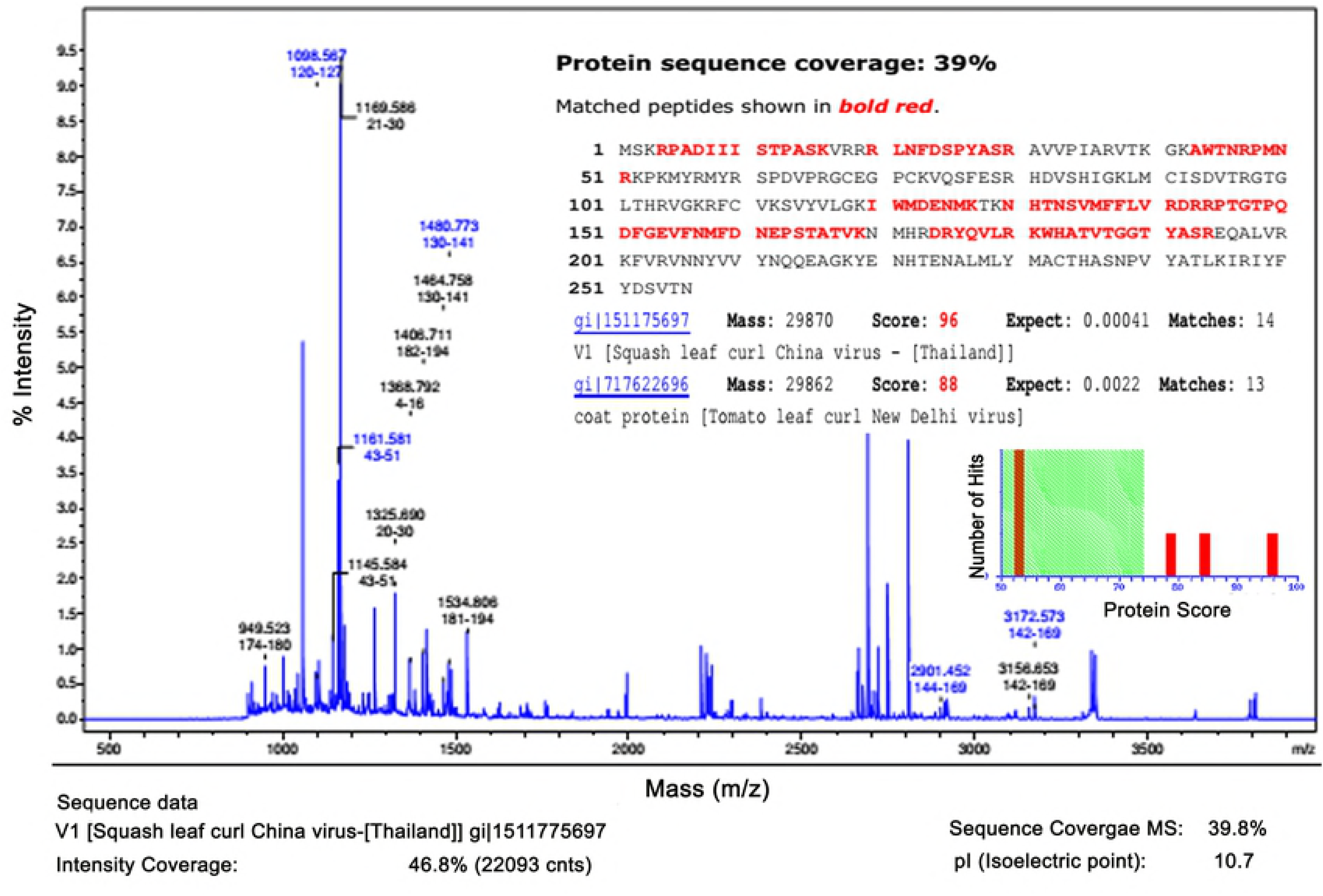

### Self-assembly of SLCCNV (native and *Pichia*-expressed CP monomers)

Early investigation of disassembly and assembly of native SLCCNV particles through electron microscopy, we found that there was no evidence of either particle disassembly or assembly observed in the virus samples dialyzed against various pH buffers (5 to10). In figure 3A-C Panel 2, the HRTEM micrographs clearly show that the native virus particles remained intact, with slight morphological transitions, when dialyzed against the pH 7-9 buffers. At pH below the 7, a slightly denatured structure and aggregated particles were observed, which was definitely not the disassembly of particles. Because the virus particles sizes remained same at the measurement of HRTEM bar scale. At pH 10, the native virus particles were found in aggregated state (Figure 3C Panel 2 HRTEM image). The data obtained from DLS also supported the evidence of electron microscopic observation which was determined that the native SLCCNV particles were intact by their size (mean ∼36nm) on the pH of 7 to 9, and they aggregate at the pH >7 and pH <9 which resulted in the increased mean hydrodynamic size of the particles. The obtained DLS data is/has been exhibited as a graph (Figure 3A-C Panel 1) and the calculated mean hydrodynamic diameter (*D*_*H*_) is plotted against the corresponding dialyzed pH buffers and then represented in a graph (Fig.S4.B, supplementary data).With the direction of obtained knowledge from self-assembly study of the native virus particles, a closer investigation of the self-assembling kinetics of *Pichia*-expressed SLCCNV-CP was performed using the same DLS and electron microscopy instruments. Here, all the HRTEM and DLS data is represented by graphs and figures (Figure 4 & 5). The electron micrographs obtained from HRTEM has clearly described the valid structural transition of SLCCNV-CP in the respective assembly pH buffer. However, we have never seen such remarkable stages of spontaneous *in vitro* virus coat protein assembly in any previous studies [74].The *Pichia*-expressed CP dialyzed against the buffer pH 5 in the negatively stained HRTEM micrographs show that the CP monomers are in a completely disassembled state, possibly being in an unfolding condition due to acidic buffer medium, which was also determined by DLS hydrodynamic size measurement (Figure 4A Panel 1-3). It was determined that there particles of an average size of ∼30 nm were present in the pH 5 buffer medium. We also obtained a negative measurement value, which meant that size undetermined particles were present in the buffer medium. When we observed an event of self assembly of the SLCCNV-CP at buffer pH 6, the results in the HRTEM micrograph were surprising. In figure 4B Panel 2, the electron micrograph shows a spectacular array made of coat protein monomers that can be a disassembling event as was clearly visualised in the electron micrograph in the 100nm bar scale. It has an average size of ∼300nm which was also confirmed by DLS(Figure 4B Panel 1). We confirm a speculation about the particular disassembling event of CP monomers at the pH of 6 after analyzing the results of buffer pH 7 further. However, at pH 7, a nanoscale structure that collectively organized and assembled by CP monomers into a cargo-like architecture was observed (Figure 4C Panel 2). In the same micrograph, a single twin icosahedral structure similar to the structure of *Begomovirus* (geminate) was also observed. The DLS data also confirmed that the self-assembled arrayed particles were present in a single population within the mean hydrodynamic size of ∼250nm (Figure 4C Panel 1). Significantly, SLCCNV-CP particles dialyzed at a assembly buffer pH of 8 on the electron micrograph buffer were detected as as intact, well dispersed and having multiple spherical-shaped (cargo-like) architectures with one geminate like structure on the electron micrograph (Figure 5A Panel 2). The DLS results also greatly correlated with the HRTEM observation and it was shown that the structures were in a highly dispersed state, with two different sizes of population ranging at ∼60nm and ∼510nm (Figure 5A Panel 1). In the SLCCNV-CP dialyzed at a assembly buffer pH 9, there was no evidence of cargo-like structures observed in this HRTEM micrograph (Figure 5B Panel 2). Also, we observed aggregated population of CP monomers in the same electron micrograph. The DLS for the same (pH 9) buffer medium also revealed that there were two populations of aggregated particles with the size of ∼100nm and ∼700nm (Figure 5B Panel 1). The CP monomers in the buffer pH at 10, small-to-large population of protein aggregates were observed in the size ranges at ∼50nm and ∼1100nm (figure 5C Panel 2). Also, the DLS data were confirmed the same, whereas 90% of protein particles were in the aggregated state at the buffer pH 10 (Figure 5C Panel 1). Overall, it was determined that the rate of self-assembly of SLCCNV-CP was greater at the pH of 8. Finally, through DLS calculated mean hydrodynamic diameters (*D*_*H*_) for SLCCNV-CP in the presence of different pH were plotted against the corresponding dialyzed pH buffers (Fig.S4.B, supplementary data).

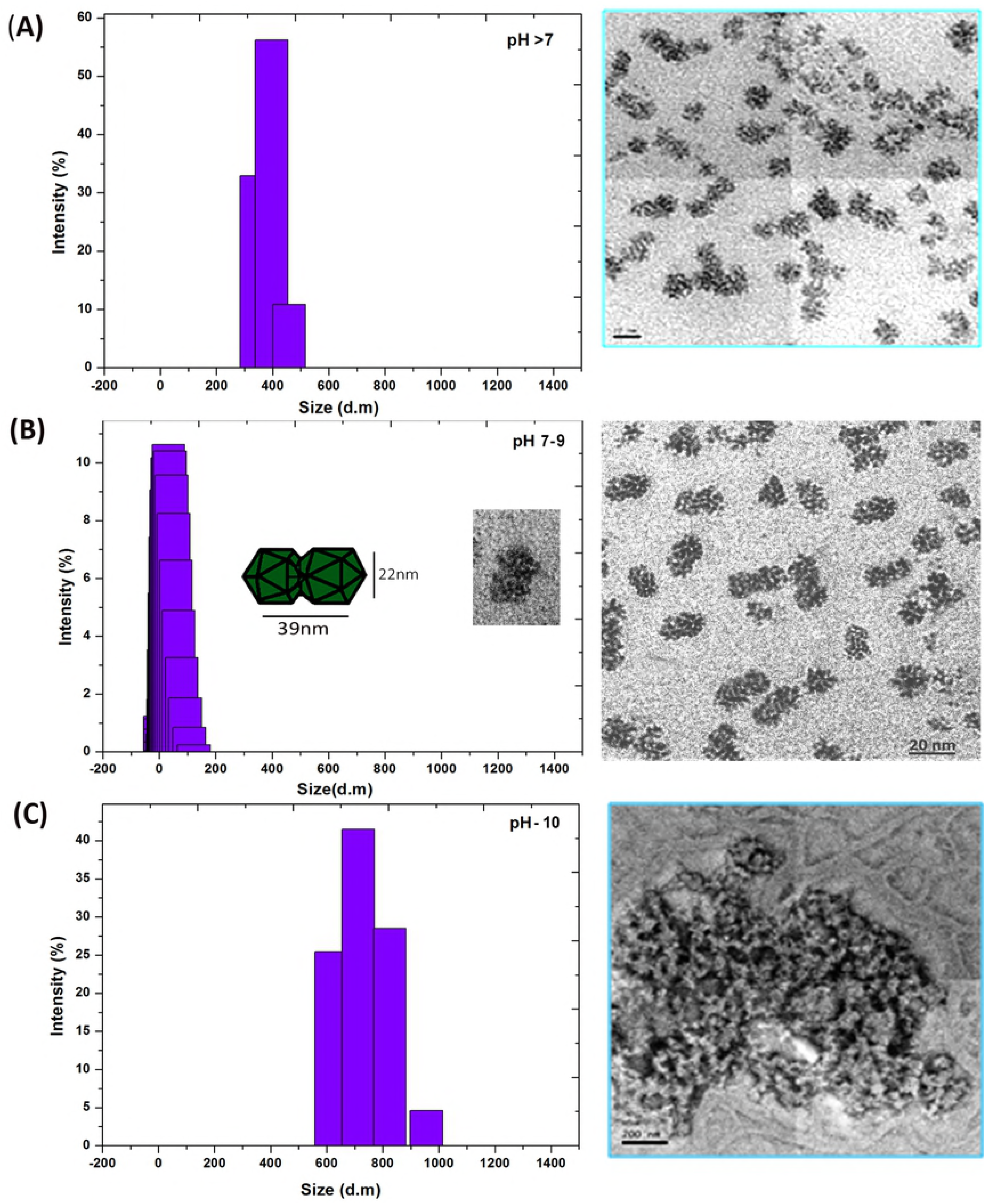

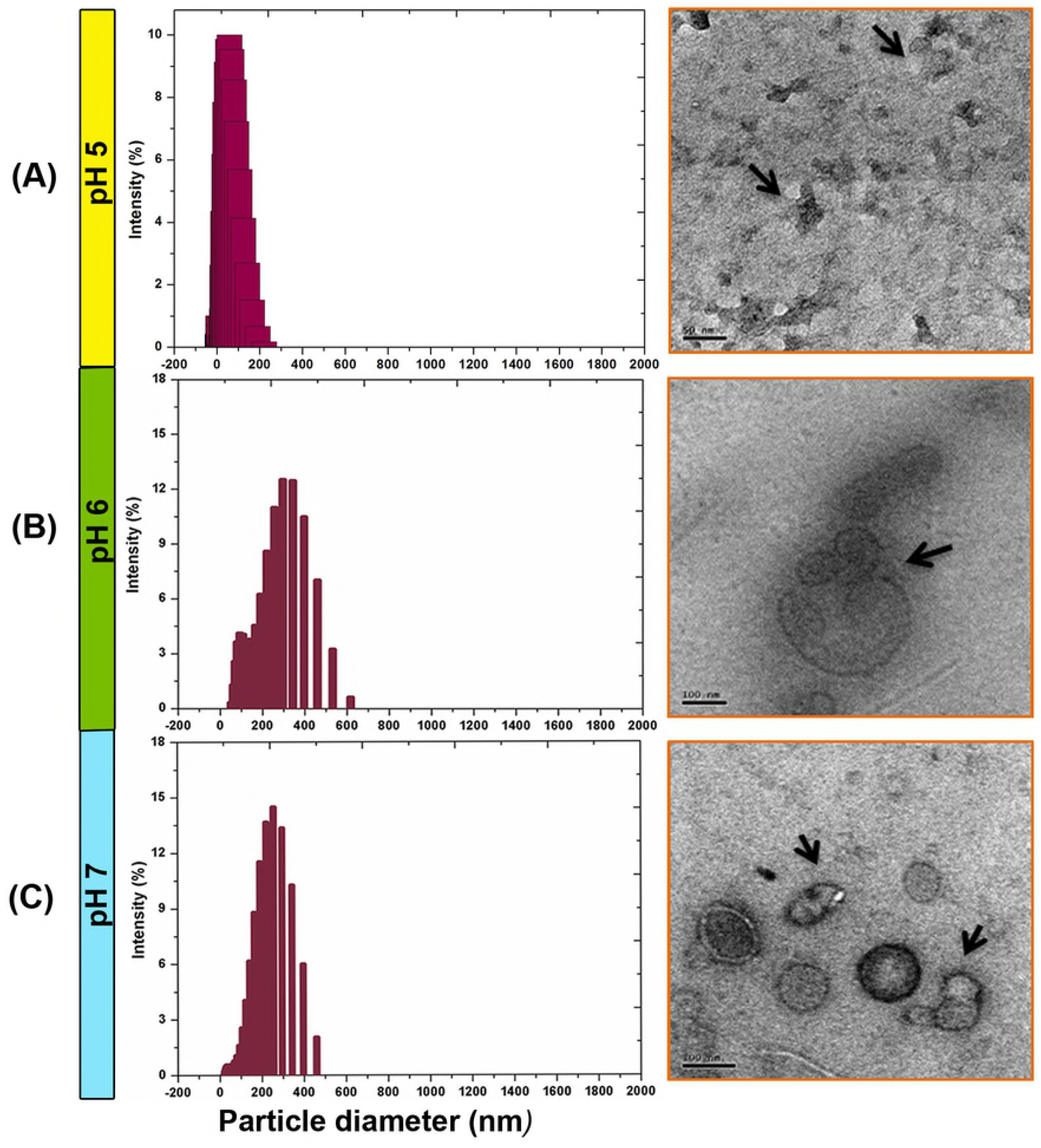

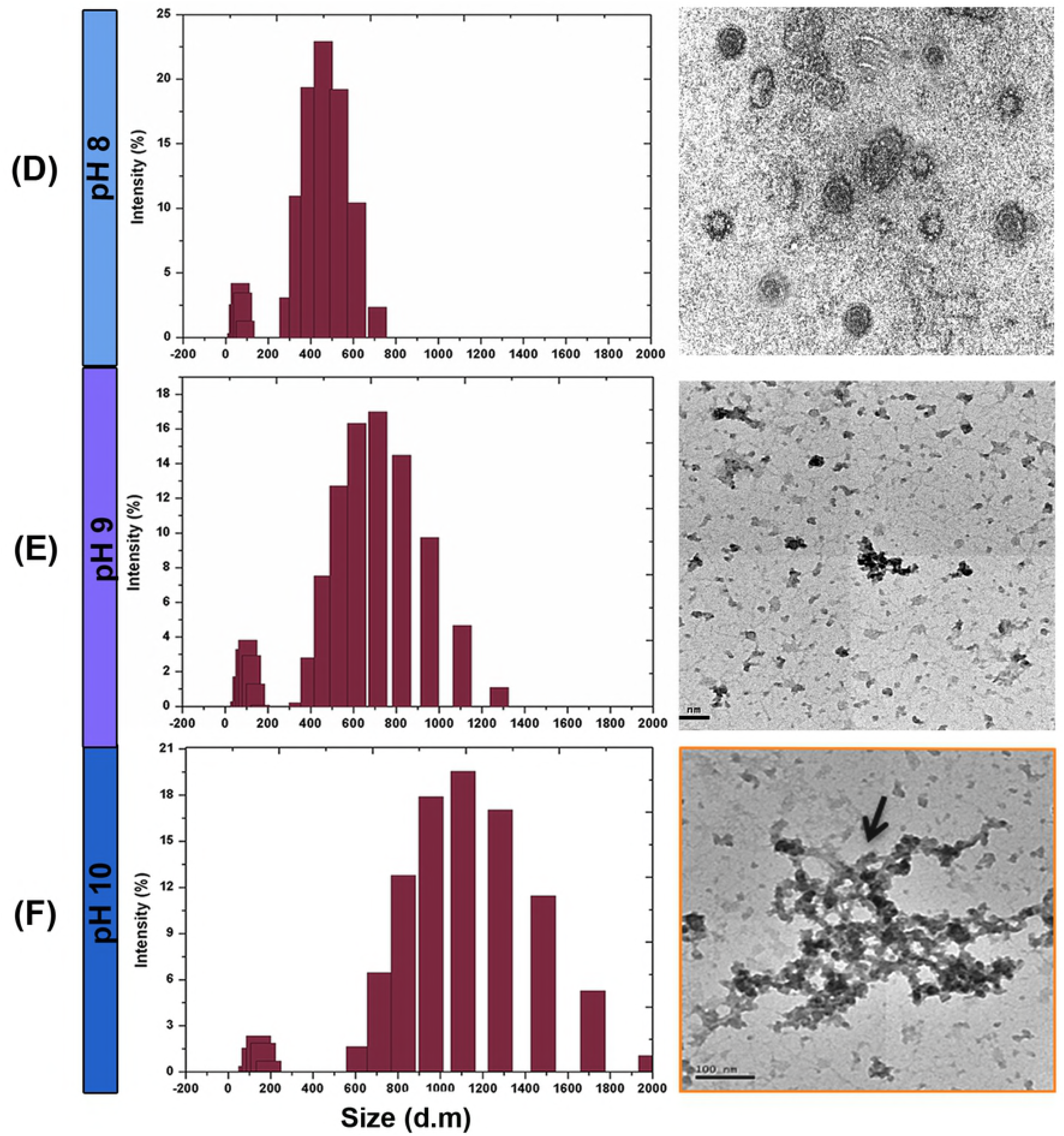

The data for the self-assembly of SLCCNV-CP at varying pH with calcium ions point out that many of the observed effects were a result of the charging of the SLCCNV-CP and their interaction with the pH medium. To interpret the results, we evaluated the capsid net charge on the basis of its peptide sequence against its respective pKa values at varying pH using an online bioinformatics tool [63]. In Fig.S4.A, supplementary data, data summation shows that the pH at which this net charge equals zero could be the theoretical isoelectric point (pI) for both SLCCNV native and expressed coat protein. Below the pI, the SLCCNV-CP has a net positive charge due to the protonation of hydrogen ions to the amino acids. The bioinformatics tool provides a logical estimation of the SLCCNV coat protein net charge in the respective buffer.

## *In silico* analysis of SLCCNV coat protein

### Modelling of monomeric protein structure in I-TASSER

The protein structure was modelled using I-TASSER, a hierarchical approach to predict the structure and function of an unknown protein from its sequence. The server generated the ten best models of protein structure, from which the best one was selected based on the C-score [64].The C-score of the predicted protein model was −2.45. The C-score of another predicted model was significantly higher and hence model 1 was chosen for further study (Figure 6A). The overall quality of the predicted protein models was evaluated using the RMSD and the TM-score, which are 0.8±0.2 and 14.2±2, respectively. The I-TASSER method was used for the prediction of protein structures with very little or no similarity, since the monomeric protein structure was not available in any of the protein data repositories [45]. The predicted protein model was validated on the SAVES PROCHECK server. The server constructed a Ramachandran plot expressing the quality of the predicted protein model (Figure 6B). The result shows that 67% of the residues are found in the favoured region, while 24% of residues are in allowed region and 9.1% of residues are in the outlier region. The 21 residues of Pro13, Arg19, Phe23, Tyr27, Thr45, Arg51, Cys68, Arg80, Asp82, Ser113, Lys119, Asp123, Glu124, Asn125, Lys127, His131, Phe159, Ser193, Phe202, Tyr219 and Asn238 were found in the outlier region. The reduced quality of the predicted protein structure was due to the amino acid residues in the outlier region. The less availability of similar protein templates resulted in the low quality of plot [55].

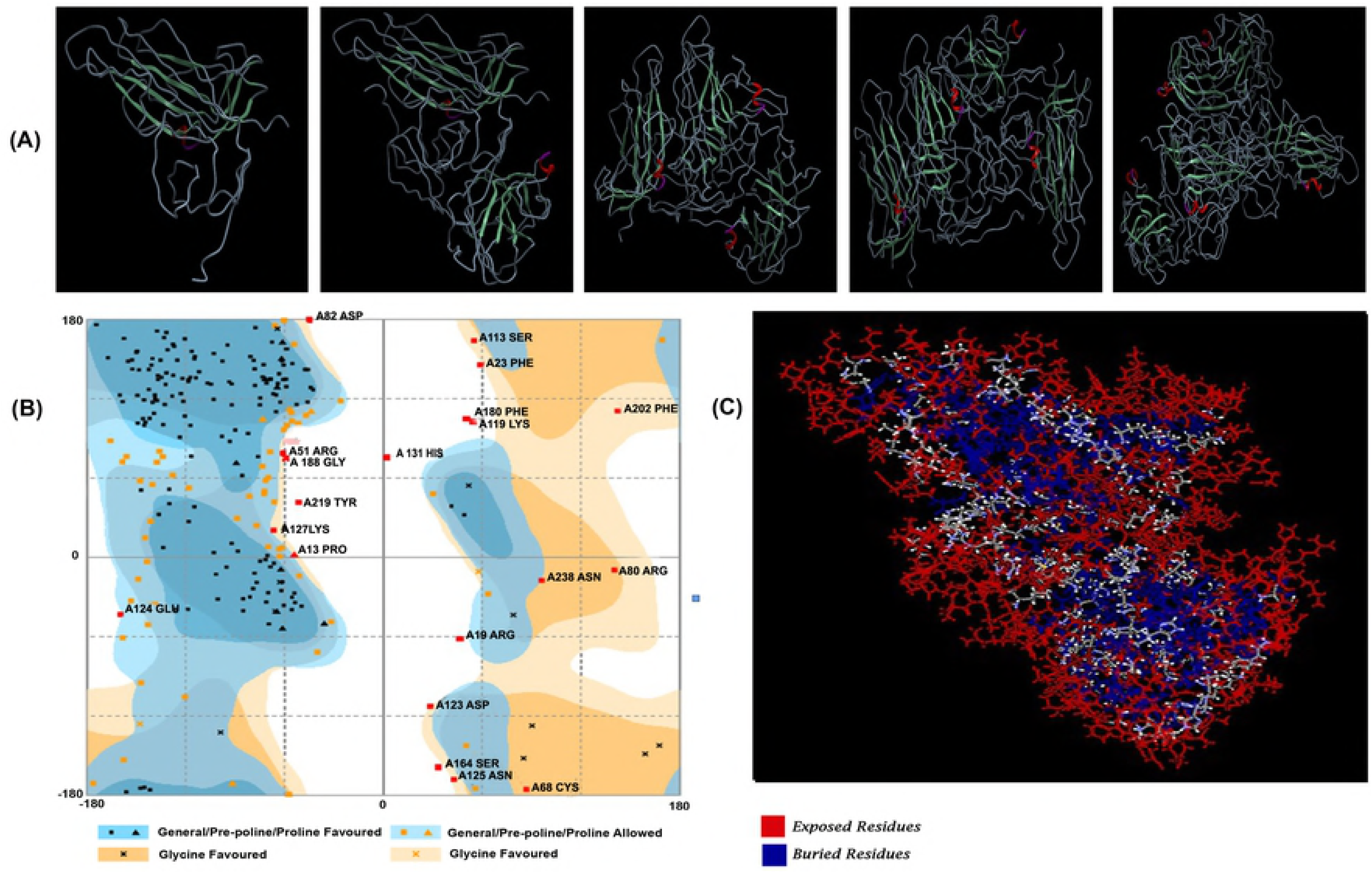

### Assembly of monomeric subunits into multimers

The SLCCNV-CP subunits were docked to assemble the monomeric units into a pentameric form of a viral capsid symmetry using a PatchDock online server. The docking method was performed with an RMSD of 4.0. The docking of monomer–monomer, dimer–monomer, trimer– monomer and tetramer–monomer was carried out in four cycles to get a pentameric structure (Figure 6A).The structure was obtained with five subunits of monomer possessing secondary structures for each subunit. The atomic contact energy (ACE) score was observed for each subunit assembly (Table 1). The docking area increased with an increase in the number of subunits, which has relatively decreased the contact energy resulting in the efficient binding of two protein structures. The atomic contact energy score is based on the number of water contacts replaced by the atomic contacts from the proteins. The atomic contact energy provides an estimation of the free energy of the protein interactions. The lower ACE configuration suggests lower free energy, which is more favourable for the formation of pentameric structure [65]. Therefore, the pentameric assembly can be achieved by monomer and dimer intermediates [66]. Using Accelrys Discovery Studio 2.5. with Connelly-type modelling, the relative solvent accessible surface areas (SASA) in the pentameric structure of coat protein could be clearly figured out. In figure 6C, it shows that the predicted solvent accessible surface, which is described in two colours (red and blue), helps in the identification of buried and exposed residues. The results shows that there are hydrophilic residues covers the entire outer region of capsomers and the hydrophobic regions are found in the deep core region.

### Analysis of binding sites and surface functional groups for bioconjugation

Three different binding sites were predicted using Molsoft ICM-Pro. The residues I14 (Isoleucine), V60 (Valine) and E78 (Glutamic acid) constitute the members of the binding site of the pentameric protein structure (Figure 7A). These functional groups are identified as being responsible for the formation of multimeric protein structure from the monomers. The surface groups were less polar compared to the accessible or the buried surface but they mostly contain charged residues. It is clear from the surface groups that non-polar interactions (van der Waals and hydrophobic) and polar interactions (hydrogen bond) present at the interfaces between the coat protein subunits are responsible for the assembly of the pentameric form into the capsomere [66]. In addition, Molsoft ICM-Pro has predicted nine different residues occupied on the outer region of the capsid protein. These residues are Lysine (K169), Tyrosine (Y176, 251), Aspartic acid (D82,151), Glutamic acid (E215,220,224), Glycine (G98,147), Asparagine (N157,170), Arginine (R91,144), Valine (V155), and Threonine (T99) (Figure 7B). These residues are the functional groups responsible for the interaction with other molecules.

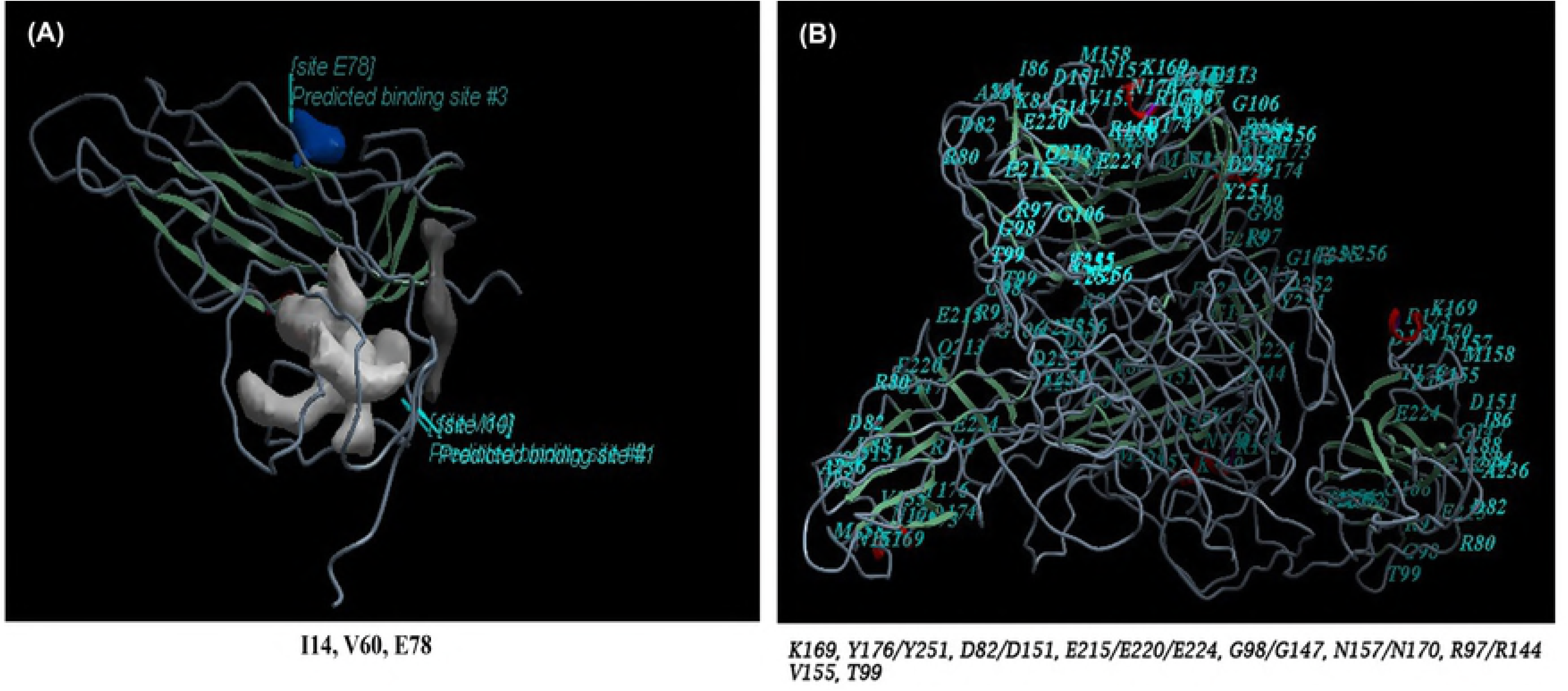

### Biocompatibility of expressed SLCCNV coat protein

The MTT assay method is widely used for testing biocompatibility and it provides the quantitative assessment of *in vitro* cell proliferation related to compatible materials. This prerequisite treatment method will be performed to test the potential compatibility issues of *Pichia* expressed SLCCNV-CP in future clinical trials. Therefore, in this *in vitro* assay, after 24 hours of incubation with SLCCNV-CP at thirty concentrations between 0.54 and 18 μg/ml, only viable cells which were capable of metabolizing MTT produced a purple coloured precipitate further analyzed by spectrophotometrically. After 24 hours of incubation, A549 lung cancer cells showed excellent viability against the SLCCNV-CP even up to the concentration of 18μg/ml (Figure 8). The IC_50_ values exceeded 18μg/ml. Significantly, the data showed that the toxicity cut-off range (70%) was reached at a protein concentration of 11μg/ml. It means that up to a concentration of 11μg/ml, SLCCNV-CP is relatively non-toxic to cells. The toxicity testing also helps to calculate the No Observed Adverse Effect Level (NOAEL) dose and is helpful in any further clinical studies [67].

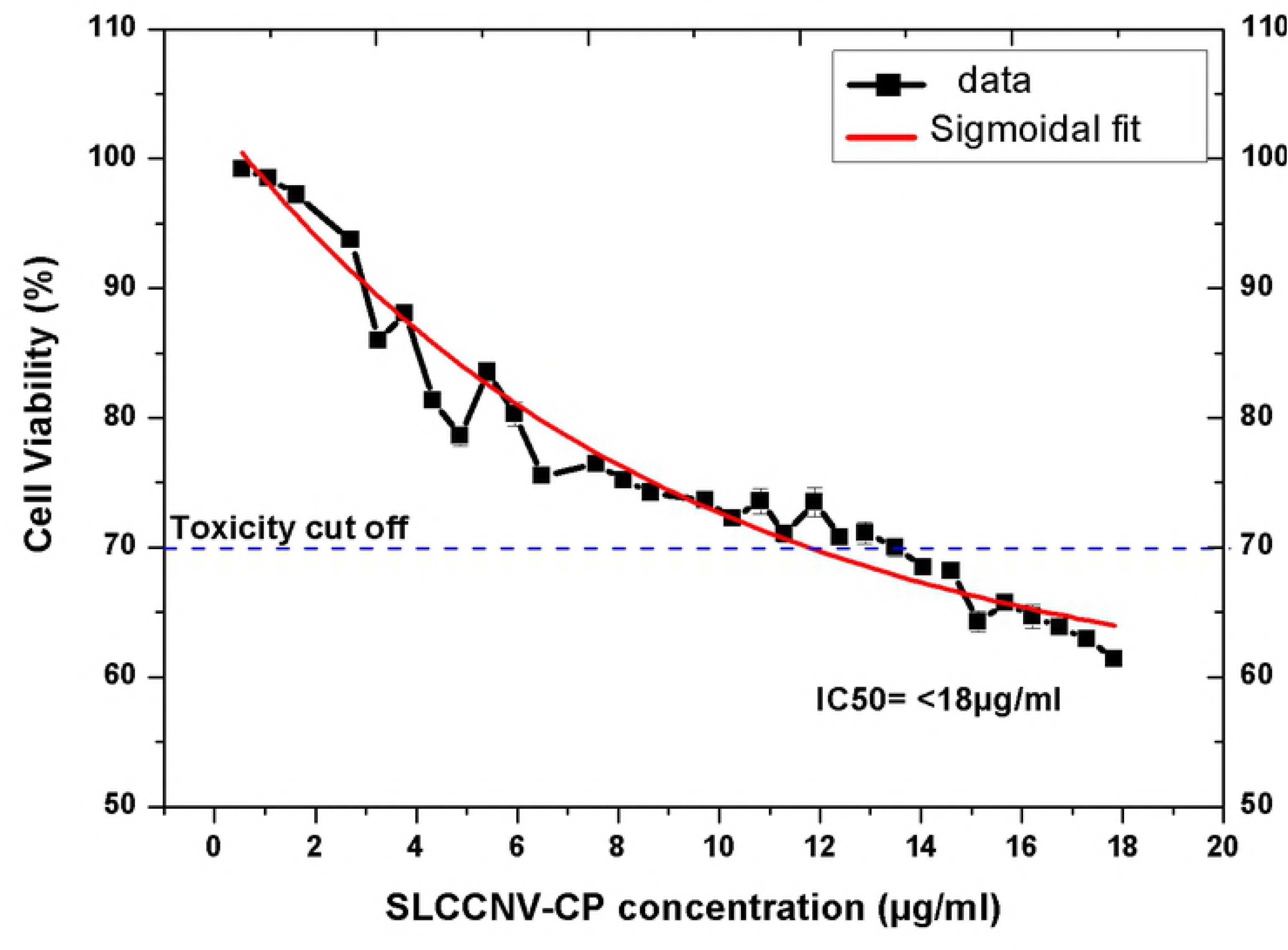

### Encapsulation efficiency of SLCCNV-CP-Nanocargo

In this approach, SLCCNV-CP monomers were mixed with MNPs at disassembly buffer (pH 6) and then dialyzed against assembly buffer (pH7). This showed prominent results after analysis through UV-Vis spectroscopy, gel electrophoresis and HRTEM. Figure 9A, the UV-Vis extinction spectra strongly represent the SLCCNV-CP monomers assembled and encapsulated around the magnetic nanoparticles core. However, the original extinction peak for unmodified MNPs changed to a low-intensity peak when encapsulated with SLCCNV-CP monomers, which was reflected by the strong difference in the extinction peak. This reversible change in the extinction spectrum clearly demonstrates that the MNPs are stabilized and might be dispersed by the particle in solution. That condition is entirely due to decreased photon absorption by SLCCNV-CP-nanocargo with MNPs. Encapsulation also confirmed at protein gel electrophoresis without staining. In Figure 9B lane 1, the visible band shows that the control unmodified MNPs were not separated. Figure 9B lane 2 shows three visible bands–one band similar to unmodified MNPs and the other two separated in the middle and bottom of the gel. Both bands resulted by the SLCCNV-CP-encapsulated MNPs, which because of the encapuslated protein helps the separation of SLCCNV-CP-MNPs electrophoretically in the gel. However, the band in the middle of the gel may resulted by few MNP particles encapsulated by SLCCNV-CP at a time, which decreased the mobility of SLCCNV-CP and kept them in the middle of the gel. The band in the bottom may be resulted by an individual particle encapsulated by SLCCNV-CP, which has increased the mobility of the SLCCNV-CP-MNPs. In figure 9C-D, HRTEM images (inset), we observed significant differences between the unmodified and SLCCNV-encapsulated MNPs. It clearly indicates that unmodified MNPs were in aggregated state while SLCCNV-CP-encapsulated MNPs were in a dispersed state. In such a case, encapsulated MNPs can be separated by a simple centrifugation method [68]. The EDX analysis also detected higher counts of C,N,O and H atoms in encapsulated MNPs, which corresponds to the atomic complexity of proteins. No such atom counts were observed in unmodified MNPs (Figure 9C-D). A scheme of SLCCNV-CP encapsulation is briefly summarized in Figure 10.

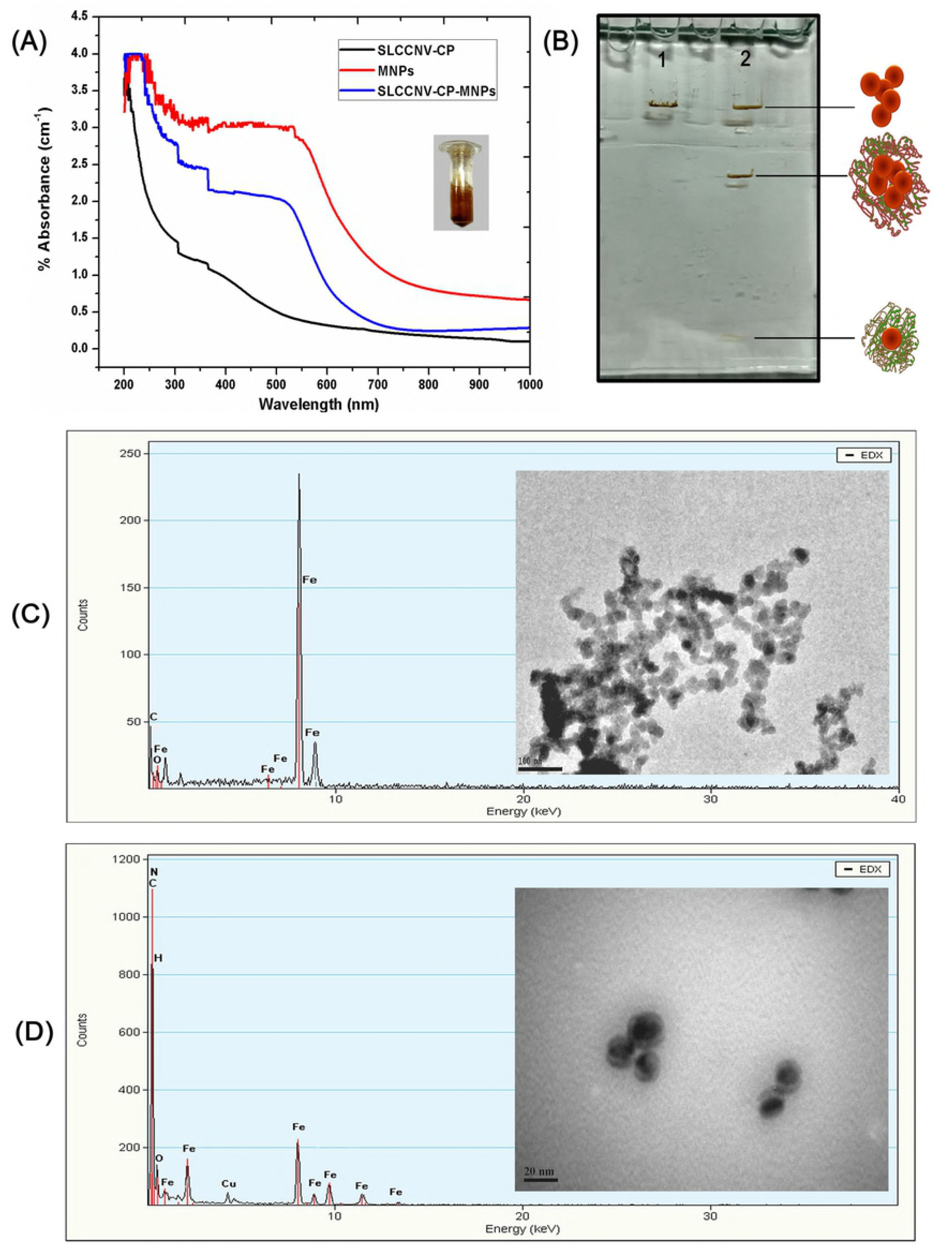

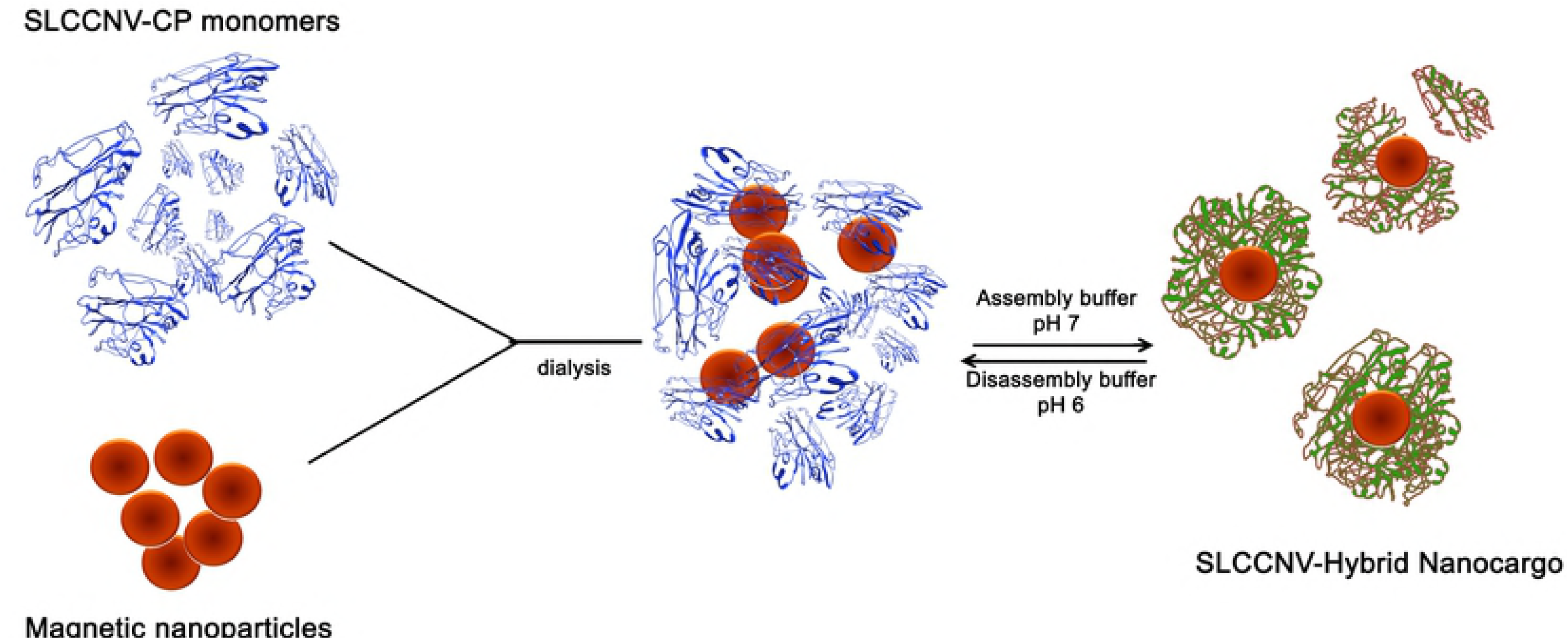

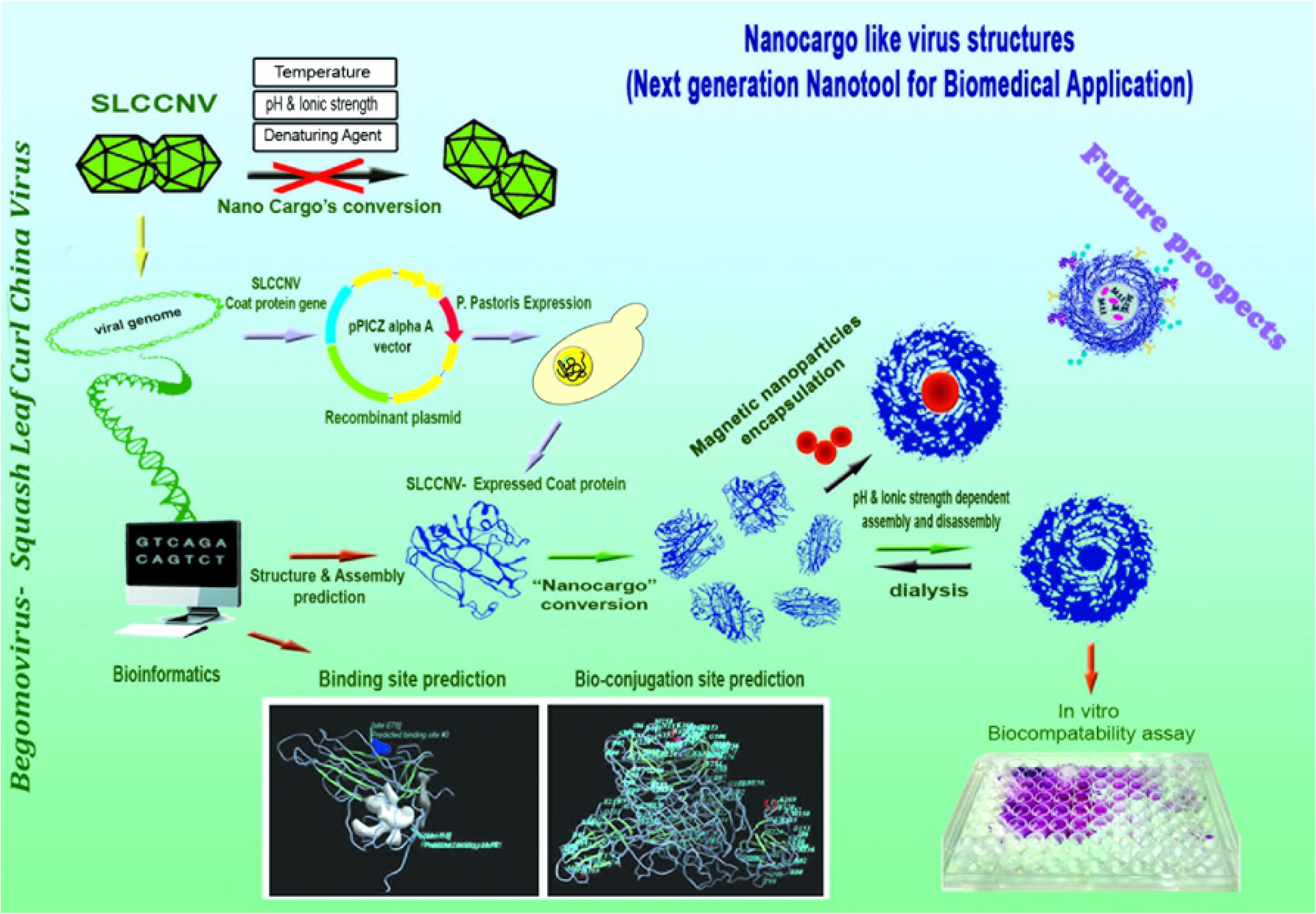

## Discussions

A key challenge in the biomedical field is the development of a smart delivery vehicle with an ability to selectively entrap therapeutic and imaging molecules and display multiple functionalities with nanoscale precision. This challenge was overcome successfully by manipulating a plant pathogenic virus to useful scaffolds in the field of biomedical nanotechnology. Here, we chose to study the self-assembling properties of *Squash leaf curl China virus* (SLCCNV) of both native whole virus and *Pichia*-expressed virus coat protein monomers. For the first time ever, complete SLCCNV-CP monomeric identical subunits of *Squash leaf curl China virus* (SLCCNV) were successfully expressed through a heterologous expression system using methylotrophic yeast *Pichia pastoris*. In our study, the usual problem associated with prokaryotic expression at post-translational modification for virus proteins (insoluble proteins) was overcome by *Pichia* expression system[19, 20]. By using pPICZαA vector specific to the secretion expression, a protein of our interest was easily isolated from the culture medium by simple centrifugation at 10,000rpm for ten minutes. As a result of these, it was easier to purify the coat protein by ion-exchange chromatography (IEC) after ammonium sulfate precipitation (Figure S3). IEC facilitates the purification of SLCCNV-CP at their pKa values. The IEC can be switched between charged and neutral depending on the pH. It was able to purify the expressed protein without any fusion proteins which might affect the homogeneity of the virus protein. Thus, using heterologous expression yields a high titer of SLCCNV-CP (0.57mg/ml) in the end compared to ultracentrifugation-purified native or native virus proteins (0.241mg/ml) (Figure S2). The sensitivity and specificity of *Begomovirus* coat protein detection was high in the Western blot analysis using polyclonal ACMV antibody. It also meant that the expressed virus proteins were properly refolded and that they retained their antigenic properties, (antigenicity) which was confirmed by the antibody’s affinity to the epitope of the coat protein. The identification of *Pichia*-expressed SLCCNV coat protein through MALDI-TOF peptide mass fingerprinting (PMF) showed a significant similarity with other *Begomovirus* species when analyzed through Mascot database searching. It was found to be quite similar to other *Begomovirus* species like SLCCNV-Thailand strain and *Tomato leaf curl New Delhi virus* (ToLCNDV). And the database search shows that there is a maximum (96%) peptide sequence similarity with the SLCCNV-Thailand strain. MALDI-TOF data could provide valuable insights into the *Begomovirus* coat protein and its key role in pathogenesis for future structural study and comprehensive research. Significantly, we confirmed the isoelectric point (pI) of the coat protein monomer as 10 by the MALDI-TOF analysis for first the time. During SDS-PAGE, we observed a novel feature of the *Begomovirus* capsid structural integrity which has never been reported elsewhere[17]. Denaturing agents can disintegrate viral structural proteins only at a high temperature (85˚C–90˚C), which has been confirmed by denaturing gel electrophoresis and DSC analysis. This feature was also evidenced by the *in silico* analysis, that the 3D structure of coat protein monomer has shown the existence of an abundance of random coil, alpha helix and few beta strand. This ratio in the secondary structure can cause the coat protein to be thermodynamically stable at the standard protein denaturing temperature [69]. Moreover, the native virus particles were tremendously stable during dialysis against both acidic and alkali buffers, as were evidenced by HRTEM and DLS data (Figure 3A-C). It understood that only high temperatures or peptide digestive enzymes of protease (either host or virus origin) can cause the cleavage of virus capsids, making them disassociate [70]. A similar kind of stability of virus particles has been observed and reported previously [8].

This investigation of the self-assembly of *Pichia*-expressed SLCCNV-CP has provided unambiguous evidence that it has an inherent ability to self-assemble in a solution without nucleic acid. This point was revealed after examining the ultrafine structures observed on HRTEM images, which were the outcome of the *in vitro* self-assembly of numerous capsomeric proteins into a defined structure (Figure4 &5). In this experiment, the assembly pH buffers are act as a cytosolic buffer system which facilitates an *in vitro* assembly of virus coat protein similar in the natural process.

The pH must have acted as one of the fundamental driving processes for the coat protein self-assembly by the protonation and deprotonation of amino acids [71]. The data obtained from the *in silico* analysis also very much correlating with the results of pH mediated assembly. However, coat protein structure prediction reveals that the subunits consist of a larger proportion of both random coil and alpha helix, which obviously makes the SLCCNV coat protein much more affinity to a hydrogen atom (H) present in the solutions [72]. Basically, protein systems are inherently capable of forming H-bonds in their main chain and side chain amino acids. These groups can, therefore, tend to be responsive while presenting the protein at the various pH environments, and we are bound to agree with another report by Casper (1963) [73] which pinpoints the key role of pH in protein assembly. He suggests that protons are hydrogen-bonded to carboxyl-carboxylate pairs and that they prevent the normal electrostatic repulsion such groups would have on each other. If the two carboxyl groups were ionized, this repulsion could strain the subunit structure and lead to dissolution. This may account for the great sensitivity of the SLCCNV-CP assembly towards change in pH. Thus, the viral coat protein assembly process was found to be very well synchronized with pH and ionic strength [74, 29]. However, altering the pH from acidic to basic could cause a structural transition of virus coat proteins from an undetermined structure (self-assembled aggregates) into a “Nanocargo”-like structure. Besides, the DLS results were clear examples of this phenomenon of structure-associated conformational changes (Figure 4 & 5). Thereby, it can be assumed that the calcium ions also have a significant role in the assembly process next to the pH buffer. There are few studies also available to describing the impact of calcium ions in the virus assembly [20, 30, 49,50]. According to that study, calcium ions reportedly act as a chaperon to specifically bind and bring the coat protein subunits together to assemble like a virion capsid. This statement of the study also proved by our intensive study with the bioinformatics tools of PatchDock and Molsoft ICM-Pro modelling. Using bioinformatics tools we found a particular intermediate amino acid residues constituting the members of the binding site between the coat protein monomers identified as Isoleucine (I14), Valine (V60) and Glutamic acid (E78) (Figure 7A). Both Isoleucine and Valine are hydrophobic branched-chain amino acids (BCAA) having an aliphatic side chain and Glutamic acid is polar hydrophilic and negatively charged by its carboxylic acid side chain. Therefore, it assumed that the available BCAA residues primarily neutral at physiological pH should have aliphatic–aliphatic interactions which are known to be a hydrophobic interaction facilitates interaction between the monomer to a monomer [75].

These features also make capsomers are thermodynamically stable and also weak in the ionization of pH variation. The previous studies reveals the features of glutamic acid and its natural affinity on the calcium ion [42,53,76]. The calcium ion affinity on the viral coat protein assembly also discussed by few earlier reports [77,59]. According to their study, the *Satellite tobacco necrosis virus* (STNV) was able to assemble in the presence of calcium ions at interfaces between the coat protein subunits. They infer/concluded that incorporation of calcium ions into aspartate and glutamate residues contributed to coordinate the STNV capsid assembly in a proper symmetry. Also some studies cited that incorporation of calcium ions into plant virus capsid is typically observed and thought to be involved in viral assembly [29,30,49,59]. Now it has become clear that BCAA residues (I14, V60) and glutamic acid (E78) residues are the intermediates that serve as a principal inter-monomer contact, and these contacts are stabilized by pH and calcium ions. Those intermediate interactions presumably can assemble the monomeric form of tertiary structures with further assembly around the existing protein particle that acts as nucleation point, which resulted in high structural organization of quaternary structure of pentamer that must possess a buried hydrophobic core [78]. Thereby, under physiological conditions similar to the intracellular environment, SLCCNV-CP monomers randomly moved and bound one another by their complementary binding sites, resulting in a novel ‘‘Nanocargo’’-like structure without the aid of an infectious nucleic acid. From the SASA docking results, it was established that there were no hydrophobic residues exposed in the outer regions of the pentameric structure predicted (Figure 6C). Therefore, solvent accessible hydrophilic residues of possibly positively charged residues of lysine, arginine and histidine might be coordinate with negatively charged nucleic acids can facilitate an error-free assembly of a capsid structure [79]. However, the self-assembly and resultant particle morphology is completely reversible upon variation of pH and the number of capsid proteins present in the reaction solution. Any increase in concentration can cause further aggregation around the existing protein particle aggregation that acts as nucleation point, which contributes to the cooperation effect [80].

Certainly, there is much demand for ingeniously self-assembled protein as a cargo material for usage in targeted therapeutic delivery [4]. Moreover, protein surface bound amino acid residues are most suitable as bioconjugation sites than chemically modified ones [81].Our self-assembled SLCCNV-CP-Nanocargo are composed of many identical copies of coat protein which consist of amino acid side chains found to be most suitable for bioconjugation than any others previously reported [8].These were predicted by the same Molsoft ICM-Pro bioinformatics tool which has predicted nine potential conjugation sites available on the exterior surface of SLCCNV coat protein monomer such as Lysine, Tyrosine, Aspartic acid, Glutamic acid, Glycine, Asparagine, Arginine, Valine and Threonine (Figure 7B) [81-82]. No free Cysteine side chains predicted which is one of the popular site for the bioconjugation methodology. The predicted available side chains, particularly amine group containing Lys, carboxylate group containing Asp then Glu, aromatic group containing Tyr are the most common bioconjugation residues utilized for the conjugation chemistry for a stable covalent link between molecules [8]. Moreover, they are the groups predominantly available on the surface of the pentameric coat protein structure. The remaining amino acids Gly, Asp, Arg, Val, and Thr are used as alternatives and specifically to tether the molecules via ionic and hydrophobic interactions with chemical intermediates. These are the residues of functional groups responsible for the interaction of the protein with other molecules [81-82]. In this regard, SLCCNV-CP-Nanocargo can be functionally modified with any kind of molecules to become “smart hybrid materials” for potential applications. It is also possible to encapsulate any small molecules (either polar or non-polar), in a single step by SLCCNV-CP-Nanocargos [8]. Their efficiency would be directed by the concentration and the relative ratio between SLCCNV coat protein and specific molecules, making them one of the finest virus hybrid materials. An important question that we have the answer now is about the biocompatible properties of SLCCNV-”Nanocargo”. There are only few studies reported the biocompatible properties of plant virus nanoparticles previously [8]. From the cell cytotoxicity assay, it was strongly determined that using plant virus like particles for furthering any biomedical application studies would be considered safe. The purified SLCCNV-CP has proved to be a potential biocompatibility material among the existing classes of protein biomaterials and that it can be used without any modification to enhance the pharmacokinetic properties in the clinical trials. Thus, now we have disclosed all the potential answers for the possible SLCCNV coat protein assembly both experimentally and computationally.

Finally, the capability of self-assembled SLCCNV-CP as “Nanocargo” was determined by using magnetic nanoparticles (MNPs) as a core to encapsulate coat protein monomers. Under optimal assembly (pH 7.0 - 8.0) conditions, encapsulation occurs spontaneously by mixing the protein subunits with cargo particles (MNPs) at a detectable level by UV-Vis absorbance, SDS-PAGE, HRTEM and EDX (see figure 16). The absorbance peak variation between free MNPs and coat protein-encapsulated MNPs clearly denotes the possible “Nanocargo” process (figure 16 A). Similarly, the principle of the separation technique of SDS-PAGE separated both free MNPs and coat protein-encapsulated MNPs by size (figure 16 B). The SLCCNV coat protein facilitates the mobility of encapsulated MNP and MNPs electrophoretically. At the HRTEM images, the fine results were observed, which correlated well with other results in this section (figure 16 C & D). This kind of encapsulation initiated via electrostatic interactions of the coat proteins with negatively charged magnetic particles results in SLCCNV-CP as an artificial Nanocargo. It is not surprising, then, that, the *in vitro* reconstitution of such virus coat protein is driven by electrostatic interactions. It is for the same reason that the virus coat protein shows an amazing ability to encapsulate not only its genetic materials but also non-native genetic materials like surface functionalized nanoparticles and other molecules. An understanding of the physical principles underlying the spontaneous encapsulation of negatively charged particles could help realize the envisaged applications in medical imaging and controlled drug release. Now, the objective of this work has helped reach the conclusion that using SLCCNV-CP-Nanocargo, encapsulation can be enabled in any other species (genomic or non-genomic) at the specific ionic strengths in the medium.

## Conclusions

Self-assembling properties of virus capsid proteins is a realistic and attractive material. The virus capsid proteins can advance the development of fabricating a robust nanotools for the nanotechnology application. Hence, we began to manipulate one of the well-known destructive plant pathogenic viruses – *Squash leaf curl China virus* (SLCCNV) – towards self-assembled “Nanocargo”-like virus structures. In this experiment, *Pichia* expressed SLCCNV-CP showed a high affinity towards pH along with calcium ions, which spontaneously facilitated the reformation of monomers to constraint “Nanocargo”-like structures. The resultant SLCCNV-CP-Nanocargo’s are stable under a pH similar to physiologic pH, and it can be disassemble simply by changing the pH to acidic. Using this pH-responsive assembly/disassembly mechanism, the magnetic nanoparticles (MNPs) were successfully entrapped into “SLCCNV-CP-Nanocargo” materials. This pH-responsive gating mechanism is a feasible way to accommodate various kinds of therapeutic and diagnostic molecules within the “SLCCNV-CP-Nanocargo”, which will address them as a next generation nanotool with promising biomedical applications. In fact, pH-responsive materials are in demand in the area of target therapeutic delivery [83]. For example, this type of pH-responsive reversible materials allows triggered release of therapeutic materials to an acidic environment at the tumour site [84]. Moreover, through a bioinformatics study, we sufficiently gained some realistic information about coat protein assembly, which correlates highly with the results obtained from *the in vitro* experiment. Also, another phase of *in silico* analysis deciphered the potential bioconjugation sites available in the SLCCNV-CP which could be work favourably to tether any functional moiety to deliver at the tissue target site. Significantly, it was proved that “SLCCNV-CP-Nanocargo” has no cytotoxic effects in maximum concentrations to mammalian cancer cell lines which can manifest “SLCCNV-CP-Nanocargo” as a potential biocompatibility material. Even though there are hundreds of virus-like particles studied, our extensive studies of the self-assembly of SLCCNV-coat protein show unique features which are very new to viral nanotechnology and which will also throw much light on smart delivery vehicle research in the biomedical field.

## Abbreviations

SLCCNV: Squash leaf curl China virus
CP: Coat protein
VNPs: Viral nanoparticles
VLPs: Virus-like nanoparticles
HRTEM: High resolution transmission electron microscopy
DLS: Dynamic light scattering
DSC: Differential scanning calorimetry
MTT: [3-(4,5-Dimethylthiazol-2,5-diphenyltetrazolium bromide]
IEC: Ion exchange chromatography
MALDI-TOF: Matrix-assisted laser desorption ionization-time of flight
BCAA: Branched-chain amino acid
MNPs: Magnetic nanoparticles

